# The E2F1 regulon orchestrates a proliferative emergency and vascular programming in fetal endothelial progenitors exposed to GDM: a sex-stratified systems medicine approach

**DOI:** 10.64898/2026.06.26.734910

**Authors:** Muyiwa S. Adegbaju, Oluseyi Babayeju, Olanrewaju B. Morenikeji, Olusola Ojurongbe, Bolaji N. Thomas

## Abstract

Maternal Gestational Diabetes Mellitus (GDM) and obesity are major drivers of the Developmental Origins of Health and Disease (DOHaD), predisposing offspring to premature cardiovascular disease. However, the specific molecular pathways that program this sex-specific vascular risk remain poorly defined due to the cellular complexity of the placenta. We sought to identify the primary regulatory engines of fetal vascular programming in a sex-stratified neonatal cohort. We analyzed purified neonatal Endothelial Colony Forming Cells (ECFCs)—the fundamental progenitors of the fetal vasculature—from pregnancies complicated by GDM and pre-pregnancy obesity. Using a sex-stratified regulatory inference framework, we decoupled the priming effects of obesity from the acute transcriptomic insult of GDM. Our findings reveal a profound functional asymmetry in fetal vascular adaptation. While male progenitors maintain metabolic resilience through *AKT3*-mediated buffering, the female fetal-placental interface undergoes a systemic proliferative emergency. This maladaptive state is driven by a massive unshackling of the E2F1-regulon (NES = 16.86), triggered by a maternal-fetal surge in CDK/MAPK signaling. This female-specific program prioritizes unscheduled cell-cycle progression at the metabolic expense of angiogenic maturation and innate immune surveillance. GDM imposes a sex-specific epigenetic scar on female fetal endothelial progenitors, characterized by a quantity-over-quality trade-off in vascular development. This identification of the E2F1-pathway as a driver of fetal vascular exhaustion provides a mechanistic basis for the increased cardiovascular vulnerability in female offspring and identifies the cell cycle as a potential therapeutic target for mitigating the long-term sequelae of GDM.

## Introduction

Gestational Diabetes Mellitus (GDM), frequently compounded by pre-pregnancy maternal obesity, represents a critical intrauterine metabolic insult that programs lifelong cardiovascular and reproductive risk in offspring [1]. Under the paradigm of fuel-mediated teratogenesis, maternal hyperglycemia and hyperinsulinemia induce permanent structural and functional alterations within the feto-placental unit during sensitive windows of organogenesis [2]. While the shared intrauterine environment suggests a monolithic metabolic burden, the feto-placental response is profoundly sexually dimorphic. Current evidence indicates that female progeny exhibit higher transcriptomic reactivity, suggesting a more complex—and potentially more vulnerable—regulatory landscape compared to males [3,4].

A fundamental challenge in defining these trajectories has been the inherent cellular heterogeneity of the placenta. Traditional bulk-tissue transcriptomic analyses aggregate diverse cell lineages, inadvertently masking cell-specific regulatory hubs through signal dilution.(5) Consequently, investigating purified neonatal cord blood-derived Endothelial Colony Forming Cells (ECFCs)—the fundamental progenitors of the fetal vascular tree—is essential to isolate the primary transcriptomic insult to the fetal-placental vasculature [6]. Recent phenotypic studies utilizing these purified progenitors have demonstrated that GDM and maternal weight gain significantly impair functional capacity, resulting in reduced self-renewal potential and delayed wound-healing kinetics [7]. Despite these observed phenotypic failures, the underlying molecular master regulators remain poorly defined. We hypothesized that the functional collapse observed in GDM-exposed ECFCs is driven by a sex-specific transcriptional hijacking that uncouples cellular mitogenesis from functional morphogenesis. To resolve this, a high-resolution approach is required to move beyond simple gene-level changes and reconstruct the underlying regulatory engines governing this divergent remodeling.

In this study, we utilized a sex-stratified regulatory inference framework and purified neonatal ECFCs to define the divergent molecular trajectories of GDM-exposed fetal progenitors. Here, we describe a fundamental functional asymmetry: whereas the male feto-placental interface preserves adaptive resilience through *AKT3*-mediated metabolic buffering, the female cohort enters a maladaptive proliferative emergency. We identify the E2F1-regulon as the master orchestrator of this female-specific response. We demonstrate that this molecular bypass induces a transcriptional collapse of the homeostatic niche, providing a robust mechanistic basis for the impaired self-renewal and delayed wound healing observed phenotypically in GDM-exposed ECFCs [7]. By characterizing the E2F1-regulon as the core architect of female vascular dysfunction, we identify the cell cycle as a primary, sex-specific therapeutic target for mitigating the developmental origins of cardiovascular disease.

## Methods

### Data source and *in silico* verification of cell identity

Transcriptomic profiles were derived from neonatal Endothelial Colony Forming Cells (ECFCs; *n* = 60) originally isolated and characterized by Weiss et al. [6]. Raw sequence data were retrieved from the NCBI Sequence Read Archive (SRA) under BioProject PRJNA885556. While the primary study utilized a subset of these data (n = 35) to investigate long non-coding RNAs in the context of gestational weight gain, we performed a comprehensive re-analysis of the full 60-sample cohort. This allowed for a sex-stratified investigation of GDM-specific protein-coding signatures, which was outside the scope of the original publication.

Primary study protocols established ECFC identity through the expression of endothelial markers (CD31, vWF) and confirmed the absence of mesenchymal contaminants (CD90, SMA). Furthermore, strict adherence to high-integrity RNA standards (RNA Integrity Number, RIN > 7.0) was maintained during the initial sequencing phase. To build upon these established standards and provide an independent quantitative purity assessment of the raw FASTQ data, our pipeline utilized the expression profiles of canonical hematopoietic markers (PTPRC/CD45, CD14) as internal negative controls (Fig 1). This bioinformatic gating strategy allowed for the verification of cellular homogeneity at the transcript level across all 60 samples prior to cohort stratification. This dual approach—leveraging the peer-reviewed isolation standards of the primary study while implementing our own transcript-level verification—ensures that the identified signatures represent intrinsic vascular reprogramming.

**Fig 1:**
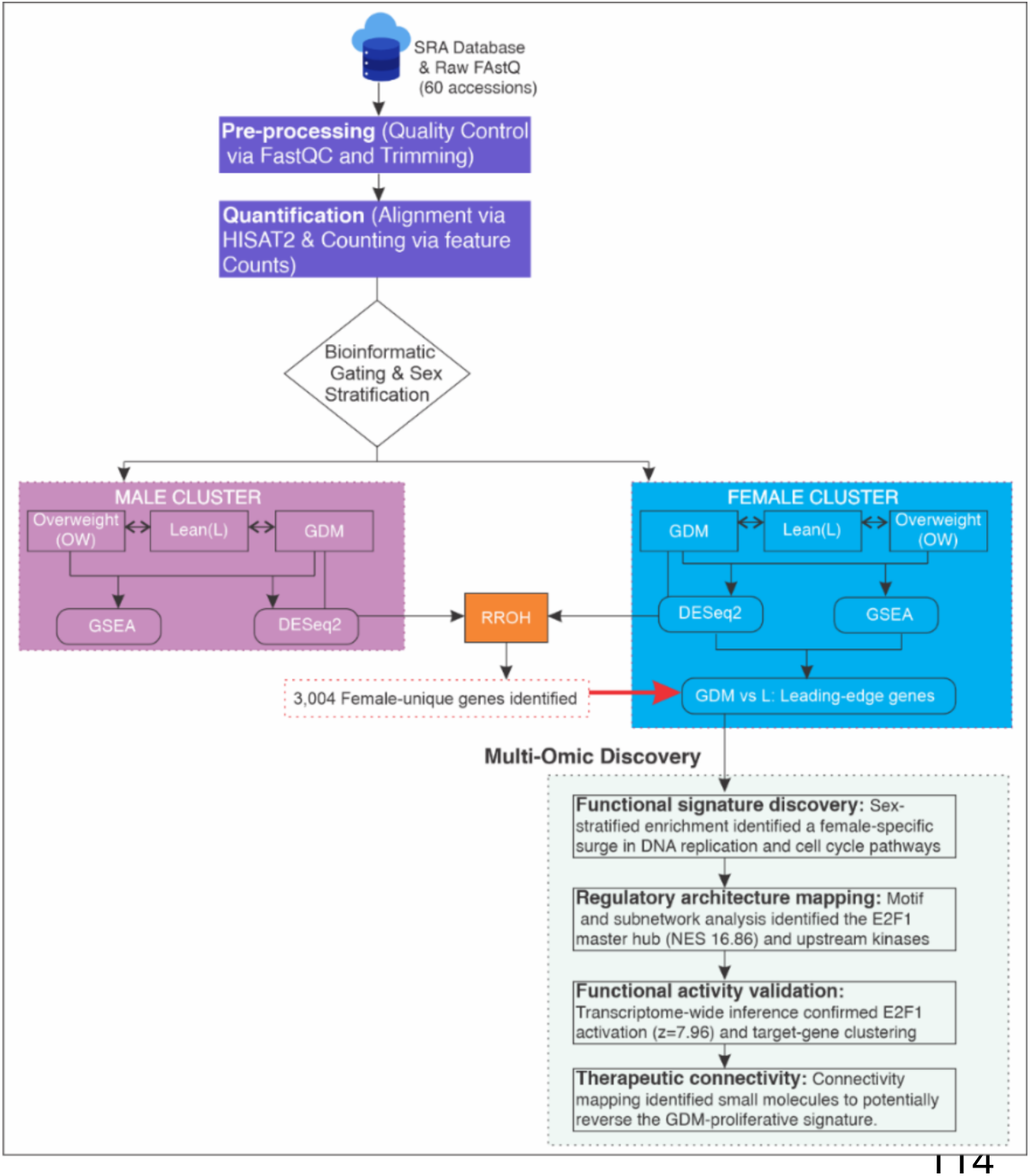
Integrated sex-stratified comparative and regulatory pipeline. Initial differential expression (DESeq2) and heatmapping compared GDM, Lean, and Overweight cohorts across both sexes. Rank-Rank Hypergeometric Overlap (RRHO) analysis of the GDM vs. Lean comparison identified a distinct signature of 3,004 female-unique genes. From this female-unique signature, GSEA Leading-Edge genes were extracted and processed via WebGestalt and GSVA, identifying a surge in DNA Replication and Cell Cycle pathways. iRegulon and X2K further mapped these to the E2F1 Master Hub (NES = 16.86) and upstream CDK1/MAPK signaling. decoupleR/CollecTRI activity inference on the full transcriptome confirmed potent E2F1 activation (z=7.96), corroborated by the clustering of validated targets (*FBN2, ANGPT2*) in an Informative Volcano Plot. Finally, the L1000FWD platform identified perturbagens capable of reversing this female-specific GDM proliferative program.

### Bioinformatics, Quality Control, and Transcriptome Alignment

Raw FASTQ files underwent initial quality assessment using FastQC (v0.11.9) [8]. To ensure high-quality alignment inputs, adapter sequences and low-quality bases were removed via Trimmomatic (v0.39) [9] using a sliding window approach (4:20). Read filtering was strictly maintained at a minimum Phred score of 20 and a minimum length of 50 bp. Cleaned paired-end reads were mapped to the human reference genome (GRCh38/hg38) using the HISAT2 (v2.2.1) splice-aware aligner [10]. Resulting alignments were converted to BAM format and coordinate-sorted via SAMtools (v1.13) [11]. To ensure the highest quantification accuracy, gene-level expression was mapped to the GENCODE (v44) primary assembly using featureCounts from the Subread package (v2.0.1) [12]. We restricted our analysis to uniquely mapped reads to eliminate signal dilution from multi-mapping artifacts, ensuring that the final count matrix represented high-confidence protein-coding transcripts.

### Statistical framework and differential expression

Differential expression analysis was performed in R (v4.3.1) using the DESeq2 (v1.40.2) package [13]. To evaluate global transcriptomic variance and confirm that maternal metabolic status and neonatal sex were the primary drivers of diversity, dimensionality reduction via Principal Component Analysis (PCA) was conducted following a Variance Stabilizing Transformation (VST) to homoscedasticize the count matrix (Fig 2). To identify the drivers of the GDM-specific phenotype, we implemented a negative binomial generalized linear model (GLM) within the DESeq2 framework, formulated as follows:

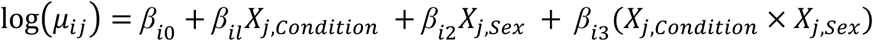

**Fig 2.**
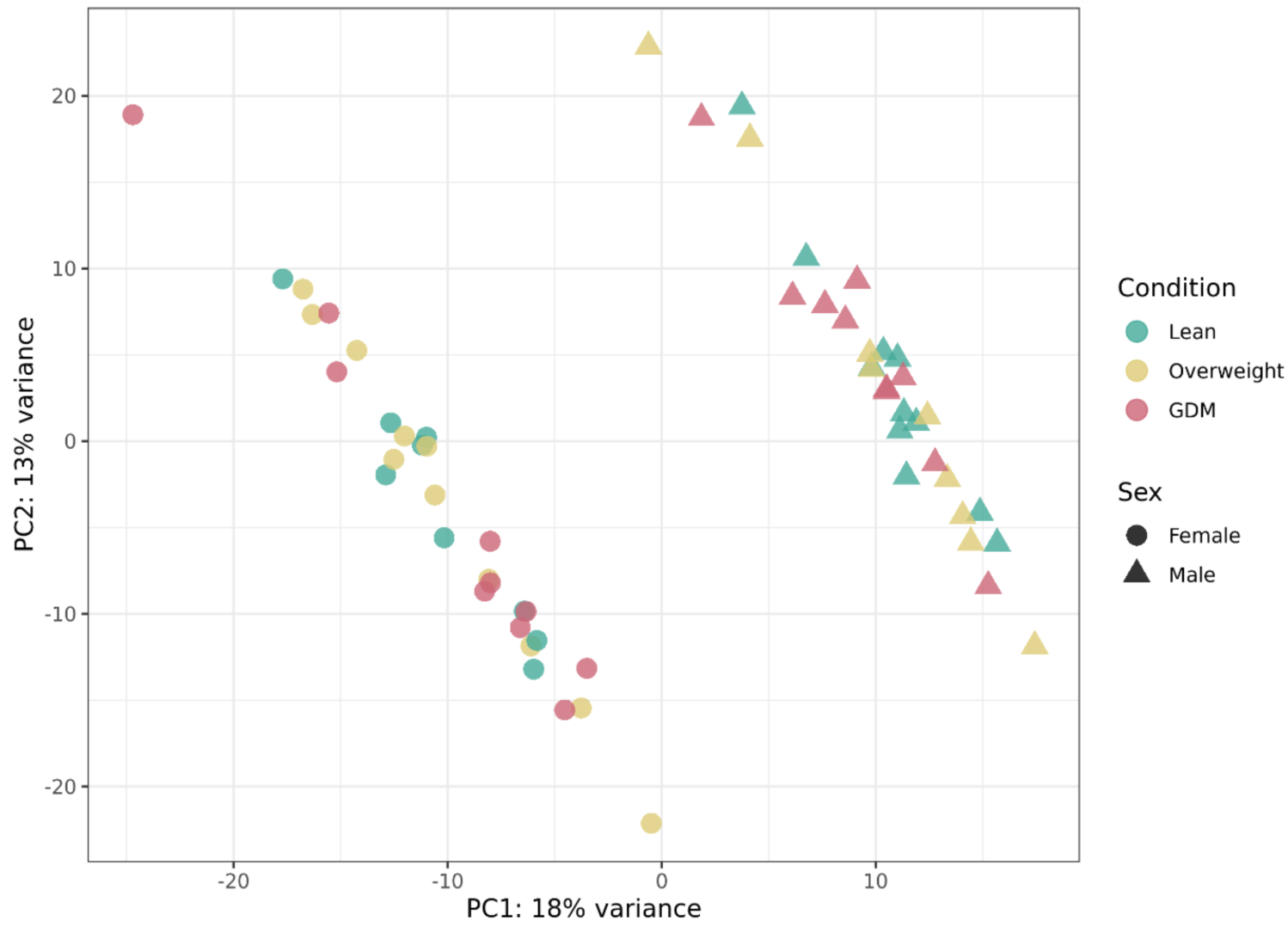
Global transcriptomic variance of neonatal ECFCs. Principal Component Analysis (PCA) of VST counts for all 60 samples. PC1 (18% variance) and PC2 (13% variance) demonstrate the primary axes of diversity. Samples are colored by maternal metabolic status (Lean, Overweight, GDM) and shaped by neonatal sex (Female: circle, Male: triangle), illustrating the baseline variance before sex-stratified analysis.

where *μ_ij_* represents the expected count for gene *i* in sample *j*, and the model evaluates the interaction between maternal condition and neonatal sex. Crucially, the analysis was stratified by sex, utilizing a balanced design (*N*=10 per group) to prevent the masking of sexually dimorphic regulatory signals. Statistical significance for pairwise contrasts (GDM vs. Lean; GDM vs. Overweight; Overweight vs. Lean) was determined using the Wald test to estimate gene-wise dispersion. To optimize the sensitivity for capturing coordinated, low-magnitude transcriptomic shifts and subtle regulatory signatures that characterize developmental programming, significant differentially expressed genes (DEGs) were defined by a nominal p-value < 0.05 and an absolute fold-change (∣log2FC∣) ≥0.58, representing a 1.5-fold difference. While unadjusted p-values were utilized to capture broader transcriptomic shifts and subtle regulatory signatures, rigorous data quality control—including Cook’s distance-based outlier replacement—was maintained to ensure the biological reliability and reproducibility of the identified master signatures.

### Rank-rank hypergeometric overlap (RRHO) analysis

To assess the global concordance and divergence of the GDM transcriptomic signatures between female and male cohorts, we performed a RRHO analysis using the RRHO R/Bioconductor package [14,15]. This threshold-free method identifies overlapping gene sets between two ranked lists without the bias introduced by arbitrary *p*-value or fold-change cutoffs. For both sex-specific datasets, genes were ranked according to their differential expression significance and directionality. A ranking metric (S) was calculated for each gene as follows:

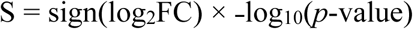

This metric ensures that the most significantly upregulated genes appear at the top of the list, while the most significantly downregulated genes appear at the bottom. To ensure statistical comparability, only the intersection of genes measured in both the female and male cohorts was utilized for the final ranking. The RRHO algorithm was executed using a step size of 200 to balance computational efficiency with high-resolution mapping of the overlap. The analysis was performed using the alternative = "enrichment" parameter to specifically identify coordinated shifts in gene expression. Statistical significance was visualized through a series of heatmaps representing the transformed *p*-values of the hypergeometric test, where the axes represent the ranks of genes in the female and male cohorts, respectively.

### Functional enrichment analysis and gene set variation analysis

To identify the biological pathways significantly overrepresented or enriched in the GDM cohorts, we performed functional enrichment analysis using the WebGestalt (v2019) online platform [16] The Gene Set Enrichment Analysis (GSEA) method was prioritized over simple Over-Representation Analysis (ORA) to include the entire ranked list of genes, preventing the loss of information from genes with subtle but coordinated expression changes. The Reactome database was utilized as the primary functional data source to map genes to well-defined biochemical reactions and signaling pathways. Pathway enrichment was determined using a minimum of 15 and a maximum of 500 genes per category to ensure biological robustness. Significance was assessed using the Normalized Enrichment Score (NES) and a False Discovery Rate (FDR) threshold of 0.05 after Benjamini-Hochberg multiple testing correction [17]. To characterize functional pathway activity at individual sample resolution, we performed Gene Set Variation Analysis (GSVA) using the GSVA R/Bioconductor package [18]. Raw transcript counts were subjected to VST normalization. Ensembl IDs were mapped to official Gene Symbols using the clusterProfiler [19] and org.Hs.eg.db libraries; for multi-mapping Ensembl IDs, expression values were aggregated using the arithmetic mean. Pathway enrichment scores were calculated using the MSigDB Hallmark (v2023.2) gene set collection using a Gaussian cumulative distribution function (kcdf) [20,21]

### Transcription factor inference and network topology analyses

To identify potential master regulators driving the female-specific GDM transcriptomic signature, we performed transcription factor (TF) motif enrichment analysis using the iRegulon plugin (v1.3) within the Cytoscape environment (v3.10.1) [22]. The leading-edge genes were analyzed using a comprehensive library of 9,713 Position Weight Matrices (PWMs) and 1,120 ChIP-seq tracks derived from ENCODE raw signals [23]. The motif search was conducted within a ranking database centered 20kb around the Transcription Start Site (TSS). A stringent NES threshold of 3.0 was applied to identify high-confidence motifs, with an Area Under the Cumulative Recovery Curve (AUC) threshold of 0.03 and a maximum FDR on motif similarity set at 0.001. To resolve the hierarchical signaling topology governing this response, the 27 female-specific leading-edge genes identified via GSEA were utilized as the discrete query set for the Expression2Kinases (X2K) pipeline [24]. TF enrichment was performed using ChEA3 [25], which compares query genes against a library of TF-target gene sets assembled from orthogonal omics datasets. To bridge the gap between inferred TFs and the upstream kinome, a comprehensive protein-protein interaction (PPI) database was deployed to identify intermediate protein interactors. The resulting subnetwork was analyzed via KEA3 to identify upstream kinases [26], which were ranked according to their Mean Rank across multiple interaction libraries (e.g., BioGRID, STRING, Kinase Library).

### Transcription factor activity inference via decoupleR

To infer the functional activity of TFs directly from the female GDM transcriptomic profile, we utilized the decoupleR R/Bioconductor package (Badia-i-Muzet et al., 2022) [27]. Unlike traditional enrichment methods, this approach estimates TF activity based on the coordinated expression of its known downstream targets (regulons) using the CollecTRI database. TF activity scores were calculated using a Univariate Linear Model (ULM), which treats the *t*-statistic from the differential expression analysis as the response variable and the regulatory influence (mode of regulation) of the TF on its targets as the predictor. A positive activity score indicates that the TF’s regulon is significantly upregulated, suggesting the TF itself is in an active state. Ensembl IDs were mapped to official HGNC Gene Symbols using the biomaRt package accessing the Ensembl BioMart database [28]. For the final visualization, genes were classified as E2F1 Targets based on their presence in the CollecTRI-validated E2F1 regulon. Differential expression significance (*p* < 0.05) and magnitude (log_2_ Fold Change) were visualized via a volcano plot generated using ggplot2 and ggrepel, with specific highlighting of the E2F1-regulated leading edge.

### Connectivity Mapping and Small-Molecule Reversal Analysis

To identify small molecules capable of reversing or mimicking the female-specific GDM transcriptomic signature, we performed connectivity mapping using the L1000FWD (L1000 Future Works Desktop) platform [29]. This analysis compares the observed disease signature against a library of over 16,000 drug-induced expression profiles from the LINCS L1000 dataset. The input signature was defined by the top 50 upregulated and top 50 downregulated genes (by *p*-value and fold change) from the female GDM differential expression analysis. The L1000FWD algorithm calculated a similarity score (ranging from -1 to +1) for each small molecule. A significant negative score (signature reversal) indicates a potential therapeutic candidate that shifts the gene expression in the opposite direction of the disease state, whereas a significant positive score identifies compounds that induce a transcriptional state analogous to GDM pathology. Compounds were further classified by their known mechanisms of action (MOA) to determine if the top reversal hits targeted the signaling nodes identified in our upstream regulatory analysis (e.g., CDK inhibitors).

## Results

### 3.1. Construction and validation of the bioinformatic pipeline

The bioinformatic workflow was initiated with a rigorous quality control (QC) assessment of the raw sequencing reads retrieved from SRA. Initial assessment via FastQC confirmed high-fidelity data, with per base sequence quality scores consistently exceeding a Phred (Q) score of 36 across the entire read length, representing a base call accuracy of >99.97%. Although a minor non-uniform base composition was observed in the initial ≈14 base pairs—a characteristic artifact of random hexamer priming during library preparation—the overall quality distribution was highly uniform across all flowcell tiles, precluding the need for aggressive trimming.

Following QC, read alignment was executed using the HISAT2 splice-aware aligner against the hg38 human reference genome. This process achieved an average overall alignment rate of approximately 98% across the entire cohort, validating both the technical quality of the source data and the precision of the mapping parameters. To mitigate potential PCR amplification biases and technical noise, Picard’s MarkDuplicates tool was employed to identify redundant molecular fragments. The analysis revealed a moderate technical duplication rate (average ∼23%), which was subsequently filtered out using SAMtools to ensure that downstream gene quantification by featureCounts considered only unique, high-confidence molecular fragments.

To ensure that the downstream immune-signaling findings were not skewed by non-endothelial transcripts, we monitored the expression of canonical hematopoietic markers within the quantified dataset. In alignment with established ECFC characterization criteria, normalized counts for *PTPRC* (CD45) and *CD14* were found to be negligible or below detection limits across all 60 samples. In contrast, pan-endothelial markers such as *PECAM1* (CD31) and *VWF* showed robust, uniform, and consistent expression across the entire cohort. These stringent preprocessing metrics and the confirmation of progenitor purity establish a robust foundation for the subsequent differential expression analysis and functional interpretation of the vascular-immune interface.

### 3.2. PCA and the statistical mandate for sex-stratified analysis

Principal Component Analysis of the ECFC transcriptomes revealed that the primary axes of variation, PC1 and PC2, accounted for 18% and 13% of the total variance, respectively (Figure 2). A discernible separation was observed between male (triangles) and female (circles) cohorts along PC1, indicating a profound baseline sexual dimorphism. Within the female cluster, GDM-exposed samples exhibited a trending divergence from lean controls along PC2, though a significant degree of overlap remained at the global level. To quantify the metabolic effect across the whole cohort, a global differential expression analysis was performed on the unified 60-sample dataset (GDM vs. Lean). This unstratified model yielded zero significantly differentially expressed genes (*padj* < 0.05), with nearly all transcriptomic features exhibiting adjusted p-values approaching 1.0 (Table S1). Together, the lack of a global metabolic signal despite visible dimorphism in the PCA indicated that the maternal environment induces highly divergent transcriptomic programs in male and female cells that cancel each other out in pooled analyses. These findings provided a rigorous statistical mandate for the sex-stratified analytical framework subsequently employed to unmask the latent, sex-specific regulatory signatures of GDM exposure.

### 3.3 Sex-stratified transcriptomic profiling and identification of metabolic signatures

Differential expression analysis (DEA) was performed to characterize the transcriptomic landscapes of GDM, Overweight, and Lean cohorts (threshold: *p* < 0.05; *FC* ≥ 1.5). Unsupervised hierarchical clustering revealed significant transcriptomic heterogeneity within treatment groups (Figure 3), though sex-stratified analysis identified distinct molecular signatures established by maternal body mass index BMI prior to GDM onset. In the female cohort, the response to pre-pregnancy obesity was characterized by the significant upregulation of *FAP* (Fibroblast Activation Protein), *SLITRK4*, and *CORIN*. Conversely, the male cohort exhibited a transcriptomic profile dominated by AKT3 upregulation and alterations in *CETP*.

**Fig 3.**
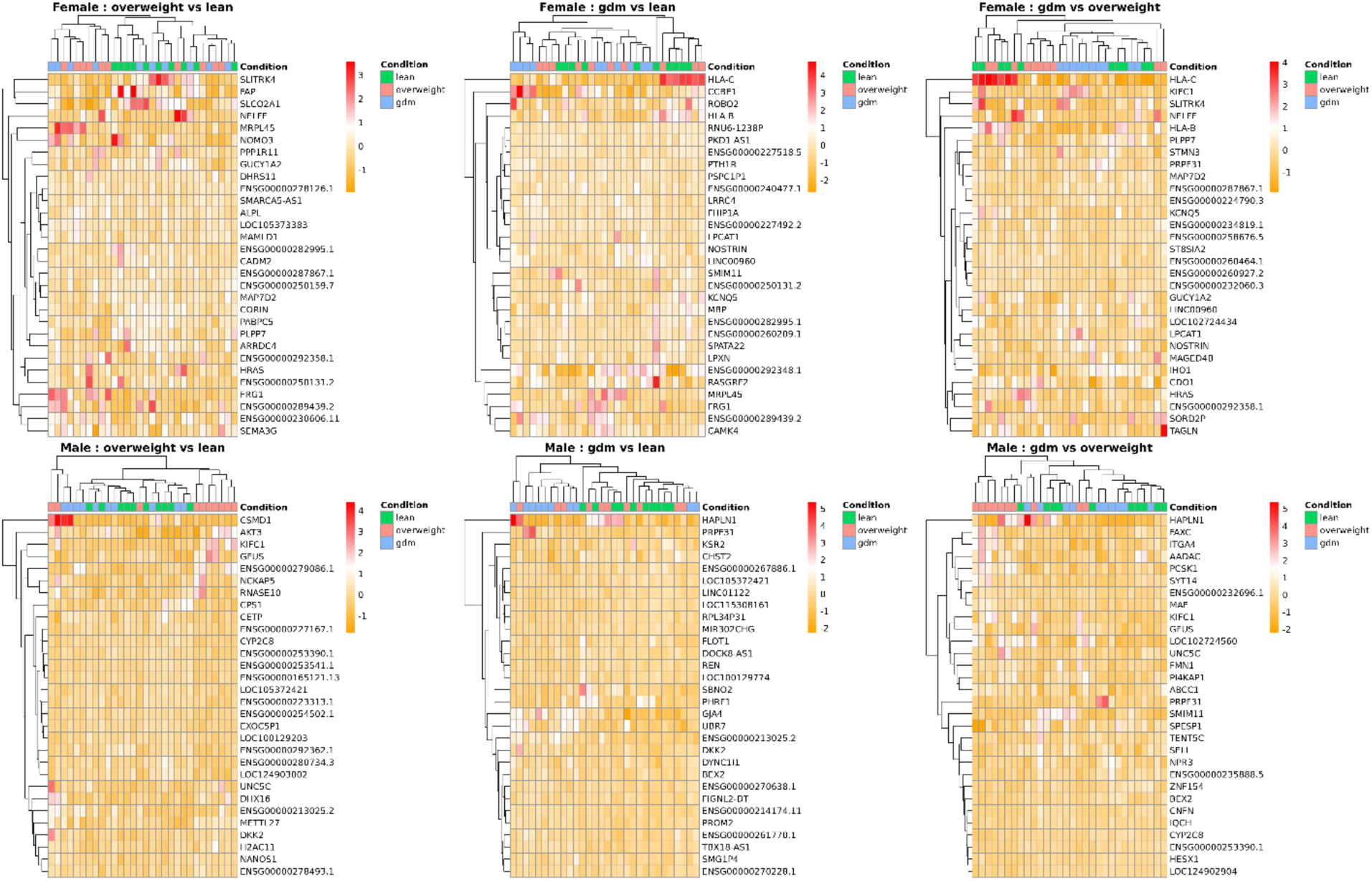
Sex-stratified transcriptomic profiling and assessment of sample heterogeneity. Multi-panel heatmaps displaying the top 30 differentially expressed genes (DEGs) across female (top row) and male (bottom row) cohorts. Comparisons are organized by baseline obesity (Overweight vs. Lean), the global impact of GDM (GDM vs. Lean), and the specific pathology of GDM isolated from obesity (GDM vs. Overweight). DEGs were identified using a nominal p-value < 0.05 and a ∣log_2_Fold Change∣ ≥ 0.58. Rows represent individual genes (mapped to HGNC symbols), and columns represent individual samples. The color gradient indicates relative expression levels, with deep red representing upregulation and orange representing downregulation (row-centered z-scores). Hierarchical clustering (dendrograms) reveals distinct molecular signatures and sample grouping based on metabolic status and fetal sex.

When comparing GDM samples directly against Overweight controls to isolate GDM-specific effects, further sexual dimorphism emerged. The female GDM-specific response was defined by the upregulation of *HLA-C*, *TAGLN*, and *KIFC1*. In contrast, the male response was characterized by the expression of *PCSK1*, *ITGA4*, and a robust, consistent upregulation of *HAPLN1* across all comparison groups. Ultimately, the GDM vs. Lean comparison confirmed divergent biological themes: the female profile was dominated by immune-modulation markers and a massive E2F1-driven cell cycle signature, while the male profile prioritized physiological homeostasis markers, including the renin-angiotensin system (*REN*).

### 3.4 Comparative functional enrichment and transcriptomic overlap mapping

Functional enrichment analysis was performed using Gene Set Enrichment Analysis (GSEA) against the Reactome pathway database to categorize the transcriptional changes across sex-stratified cohorts for all three metabolic contrasts (Figure 4). Pre-pregnancy obesity induced divergent functional profiles between the sexes, where the female cohort exhibited significant enrichment in Reactome pathways related to structural organization and growth signaling, specifically *FGFR signaling* (*FGFR1* and *FGFR4*) and *Gap junction assembly* (FDR < 0.05). In contrast, the male overweight signature was defined by a significant downregulation of *Cholesterol biosynthesis*, *Extracellular matrix organization*, and *WNT-dependent signaling*, establishing a sex-biased baseline prior to GDM onset. The subsequent transcriptomic response to GDM compared to lean controls revealed a distinct polarization of biological themes in the female cohort. Positive enrichment was localized to Reactome pathways associated with DNA replication and cell cycle progression, including *Activation of ATR in response to replication stress*, *S phase*, and *Mitotic prometaphase*, while a significant negative enrichment was identified in immune-modulatory cascades such as the *Regulation of Complement cascade*, *Interferon-gamma signaling*, and *Interleukin-10 signaling*. Notably, no biological pathways reached the adjusted significance threshold (FDR < 0.05) in the male cohort for this specific comparison, highlighting a more attenuated pathway-level response in males when measured against a lean baseline. To isolate GDM-specific effects from the confounding influence of obesity, GDM samples were compared directly to overweight controls, revealing that the female cohort maintained a consistent enrichment in Reactome cell cycle and mitotic pathways such as the *G1/S transition*. Conversely, the male cohort demonstrated a significant positive enrichment in *Cholesterol biosynthesis* and *Viral mRNA translation*. This GDM-induced upregulation of cholesterol-related genes in males stands in direct contrast to the downregulation observed during the overweight-vs-lean transition, identifying a unique and reversible metabolic shift specific to the male GDM phenotype.

**Fig 4.**
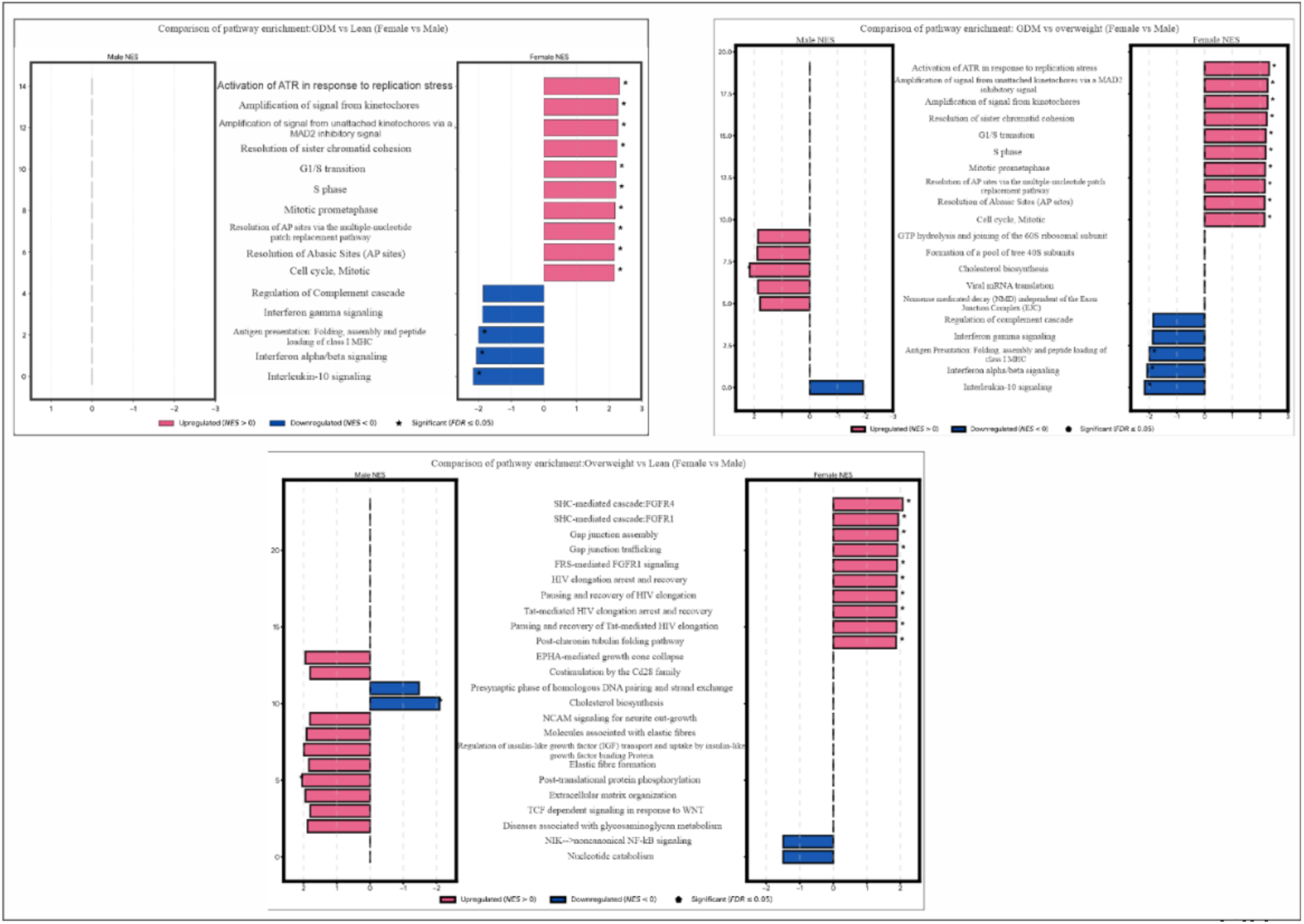
Comparative functional enrichment of neonatal ECFCs across maternal metabolic cohorts. Comparative GSEA results are presented for three primary contrasts: GDM vs. Lean (Top Left), GDM vs. Overweight (Top Right), and Overweight vs. Lean (Bottom Center). In each panel, the left y-axis represents the NES for the male cohort, and the right y-axis represents the NES for the female cohort. Pink bars denote upregulated pathways (NES > 0), while blue bars denote downregulated pathways (NES < 0). Statistical significance is indicated by an asterisk (*), denoting a FDR ≤0.05. In the female GDM cohorts, a predominant enrichment in cell cycle and DNA replication signatures is observed (e.g., Activation of ATR in response to replication stress, G1/S transition), which is conspicuously absent or less significant in the corresponding male cohorts. All pathways were mapped using the Reactome database via the WebGestalt platform.

### 3.5 Integrative convergence and divergence analysis (RRHO)

To formally quantify the transcriptomic overlap between male and female responses to GDM, we employed a dual-validation approach using Venn distribution and Rank-Rank Hypergeometric Overlap (RRHO). Initial comparison of significantly differentially expressed genes (DEGs) revealed a functional asymmetry: while 828 down-regulated genes (13.7%) were shared between the sexes, the female cohort exhibited a unique signature of 3,004 down-regulated genes. Divergence was higher in the up-regulated signatures, where only 116 genes were common to both lineages (Figure 5A). To map the global strength of this shared response, we utilized threshold-free RRHO analysis. The resulting heatmap (Figure 5B) revealed a highly partitioned signal, with a statistically significant hotspot (−log_10_(*P* -value) > 35) localized exclusively in the upper-right (UR) quadrant, representing genes strongly down-regulated in both sexes. Beyond this shared core, functional enrichment of the female-unique down-regulated signature identified a coordinated suppression of pathways involved in TGF-ꞵ signaling, ECM organization, and JAK-STAT3-mediated signaling. In contrast, the lower-left (LL) quadrant, representing shared up-regulated genes, remained a diffuse blue field, indicating a lack of concordance in the activation phase of the GDM response. While the male up-regulated profile was characterized by metabolic adjustments, the female profile showed a high-magnitude enrichment in Reactome replication-stress and DNA synthesis pathways. This divergence in the magnitude and directionality of the female genomic response provided the primary mandate for the subsequent identification of sex-specific master regulatory hubs.

**Fig 5.**
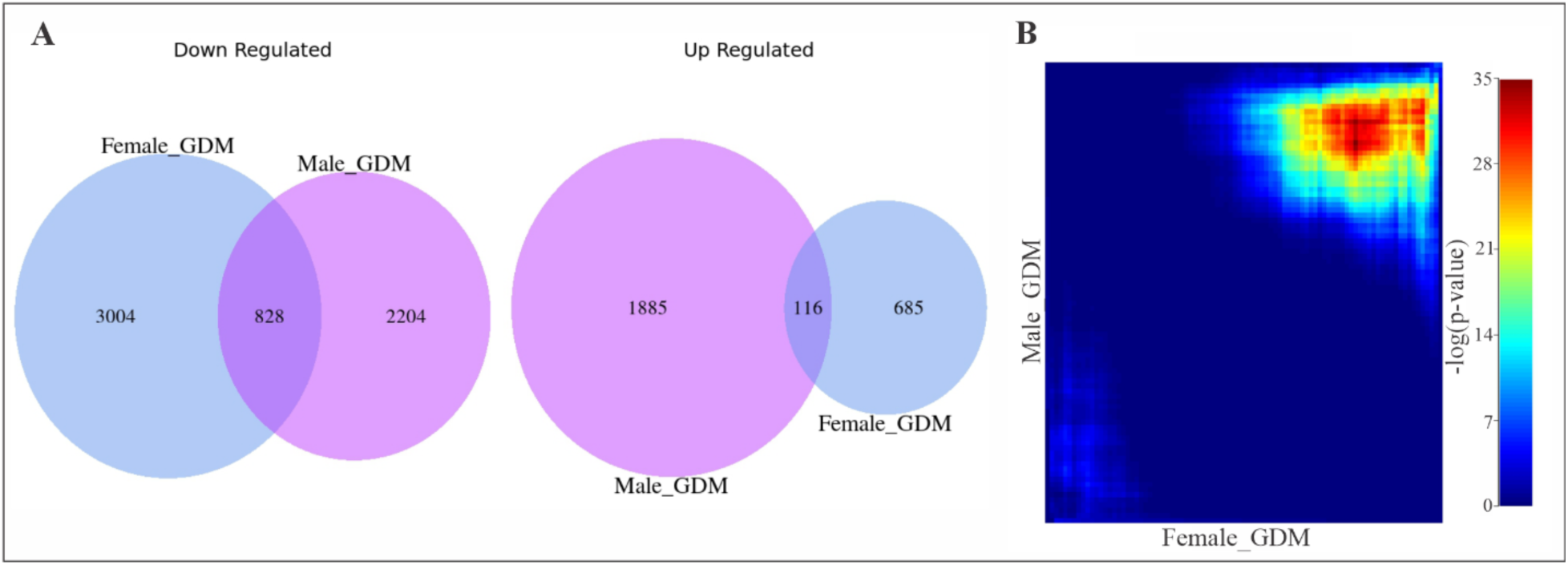
Rank-Rank Hypergeometric Overlap (RRHO) of sex-specific GDM signatures. (A) Venn diagrams of DEGs: The charts illustrate a profound divergence in transcriptomic logic, where females exhibit a massive unique down-regulatory signature (*n* = 3,004) compared to a dominant unique up-regulatory response in males (*n* = 1,885). (B) RRHO significance map: This threshold-free heatmap identifies statistically significant overlaps between female (x-axis) and male (y-axis) gene rankings. The prominent hot spot in the top-right corner reveals a highly significant shared core of up-regulated genes, including the E2F1-regulon, signaling a conserved but sex-biased proliferative emergency in response to GDM.

Although, both sexes exhibit a measurable response to GDM, the unique depth and coordination of the female transcriptomic signature—characterized by the simultaneous silencing of 3,000 genes and a high-magnitude surge in replication-stress pathways—represents the dominant regulatory event in this study. This high-resolution signal provided a clear bioinformatic mandate to focus the subsequent multi-omics discovery phase on the female lineage. By isolating this female-specific proliferative signature, we aimed to identify the core molecular architects—such as specific transcription factor hubs—responsible for orchestrating this high-magnitude genomic rewiring.

#### 3.6.1 Gene set variation analysis (GSVA) reveals female-specific pathway polarization

To estimate functional pathway activity at the individual sample level and assess the consistency of GDM-induced shifts, GSVA was performed using the MSigDB Hallmark collection (Figure 6). This transformation of the gene-level expression matrix into pathway-level enrichment scores revealed a distinct polarization within the female GDM cohort. The female GDM profile was characterized by the consistent activation of cell cycle-related signatures, with the highest positive enrichment scores observed for E2F Targets and the G2M Checkpoint. This upregulation was accompanied by significantly higher activity in Mitotic Spindle and Hedgehog Signaling pathways compared to Lean and Overweight controls. Parallel to this proliferative surge, the female GDM cohort exhibited a systemic reduction in the activity scores of pathways associated with vascular signaling and tissue organization. Specifically, IL6-JAK-STAT3 Signaling, Angiogenesis, and the Inflammatory Response showed the most robust negative enrichment. Furthermore, significant decreases were observed in the activity of Epithelial-Mesenchymal Transition (EMT) and Bile Acid Metabolism. These sample-level GSVA scores confirm that the divergence observed in the global transcriptomic analysis is a consistent feature across the individual female GDM isolations, characterized by a simultaneous increase in mitotic activity and a decrease in structural and homeostatic signaling.

**Fig 6:**
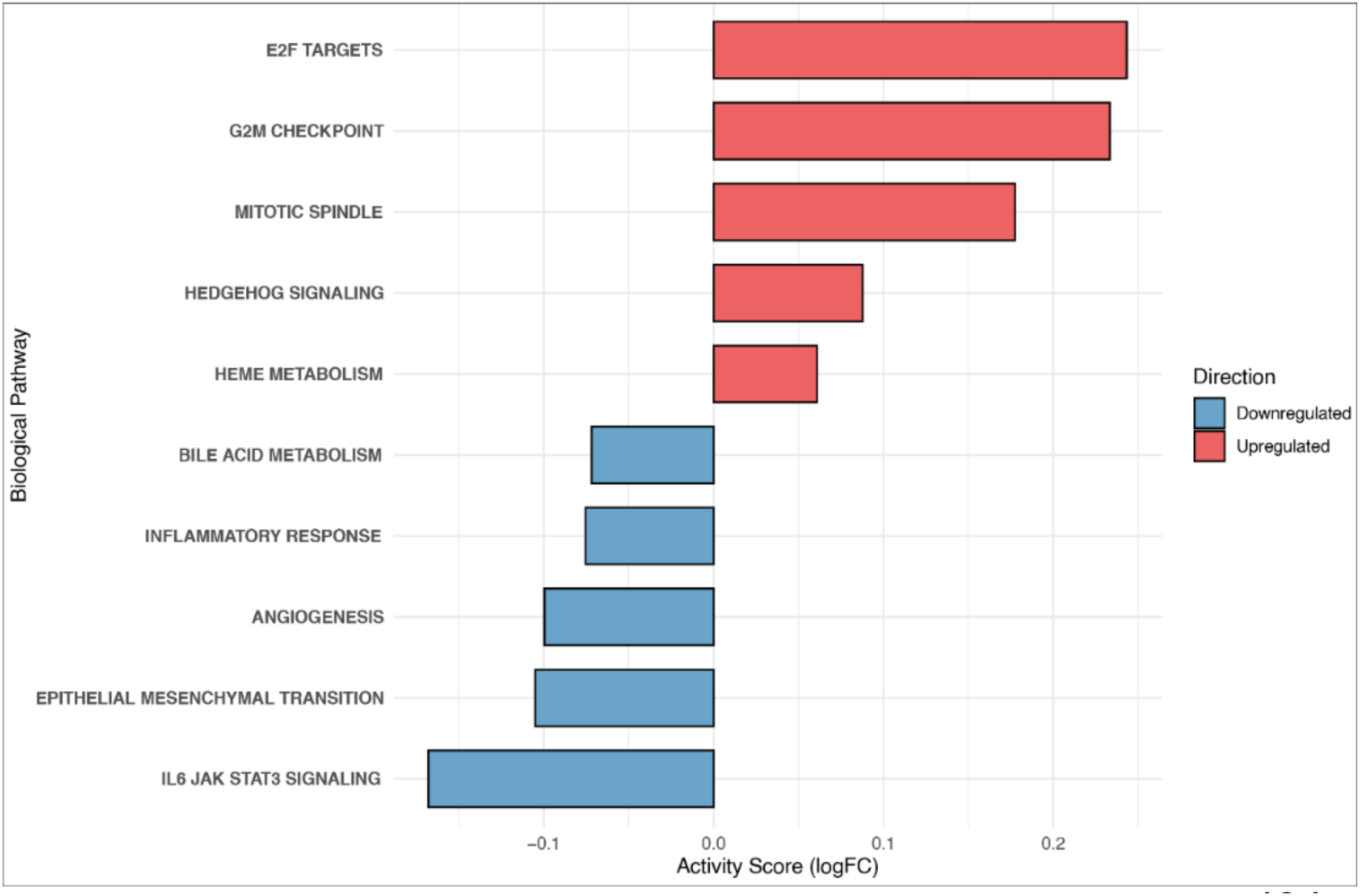
Polarization of hallmark biological states in female GDM. Bar chart displaying the differential activity scores (logFC) of MSigDB Hallmark pathways calculated via GSVA. Red bars indicate pathways with higher activity in the GDM cohort, led by E2F Targets and G2M Checkpoint regulators. Blue bars represent suppressed pathways, including IL6-JAK-STAT3 signaling and Angiogenesis. This divergent activity profile confirms a consistent shift toward a proliferative, replication-stressed phenotype in the female GDM placenta.

#### 3.6.2 Integrated upstream analysis identifies E2F1 and the CDK-kinase cascade as female-specific drivers

To identify the upstream molecular architects of the female-specific GDM response, motif-based gene set enrichment was performed using iRegulon. Analysis of the female leading-edge genes (*n* = 27) identified a regulatory signature dominated by the E2F transcription factor family (Table 1). Motif hits were grouped into functional clusters based on transcription factor homology, with the *E2F* Activation Hub exhibiting a mean Normalized Enrichment Score (NES) of 16.42. This hub, led by *E2F1*, showed high-fidelity targeting of 18 unique genes essential for the DNA pre-replication complex and the S-phase checkpoint, including the minichromosome maintenance complex (*MCM2-7*), *CDC6*, and *CDC25A*. Secondary clusters included atypical repressors (*E2F7/8*; Mean NES = 16.14) and late-cycle regulators (*E2F3/5*; Mean NES = 15.30). Notably, this *E2F*-driven regulatory architecture was absent in the male cohort.

**Table 1:** Enrichment of transcription factor binding motifs in female GDM leading-edge genes.

| Regulatory cluster | Primary candidate | Mean NES | Total unique targets | Representative target genes |
| --- | --- | --- | --- | --- |
| E2F activation hub | E2F1 / E2F4 | 16.42 | 18 | *MCM2-7, CDC6, CDC25A, CHEK1* |
| S-phase repressors | E2F7 / E2F8 | 16.14 | 17 | *MCM6, CDC6, MCM5, CDC25A* |
| Late-cycle regulators | E2F3 / E2F5 | 15.30 | 13 | *MCM3, MCM5, MCM4, CDC25A* |

To validate the functional activity of this predicted regulon, the X2K (Expression2Kinases) pipeline integrated these genes with protein-protein interaction (PPI) and kinase-substrate libraries. Inference of upstream kinases identified a coordinated signature, with *CHEK2*, *CDK2*, and *BUB1B* emerging as top-ranked regulators across multiple libraries (Figure 7A). Simultaneously, independent transcription factor analysis corroborated motif findings, with *E2F7*, *FOXM1*, and *E2F1* showing robust support across ChIP-seq and co-expression datasets (Figure 7B). The resulting X2K network illustrates a hierarchical topology where a kinase cluster (e.g., *ATR*, *ATM*, *CDK1/2*) converges upon a regulatory subnetwork dominated by *E2F1*, and *SP1*, and *MYBL2* (Figure 7C). Within this architecture, E2F1 serves as a central hub, bridging upstream mitotic signaling kinases with the downstream effector genes identified in the Reactome S-phase and DNA replication pathways. The primary mechanistic insight is revealed by the discordance between transcript levels and predicted activity (Figure 7D). Despite the high-confidence prediction of *E2F1* as a master regulator (NES = 16.86), its mRNA expression—alongside other identified TFs like *MYBL2* and *FOXM1*—remained stable (|log_2_FC| < 0.25; p > 0.05). Conversely, the metabolic trigger *INSR* and downstream transcriptional targets, including *MCM6*, *MCM2*, and *CDC6*, exhibited significant positive fold changes. This pattern—stable transcription factor expression paired with the coordinated upregulation of a validated regulon and its upstream kinases (e.g., *CDK1*, *CDK2*)—is a hallmark of post-translational activation. These data suggest that GDM-induced stress does not require *de novo* synthesis of *E2F1* but rather triggers its activity via the identified upstream kinase signaling relay.

**Fig 7:**
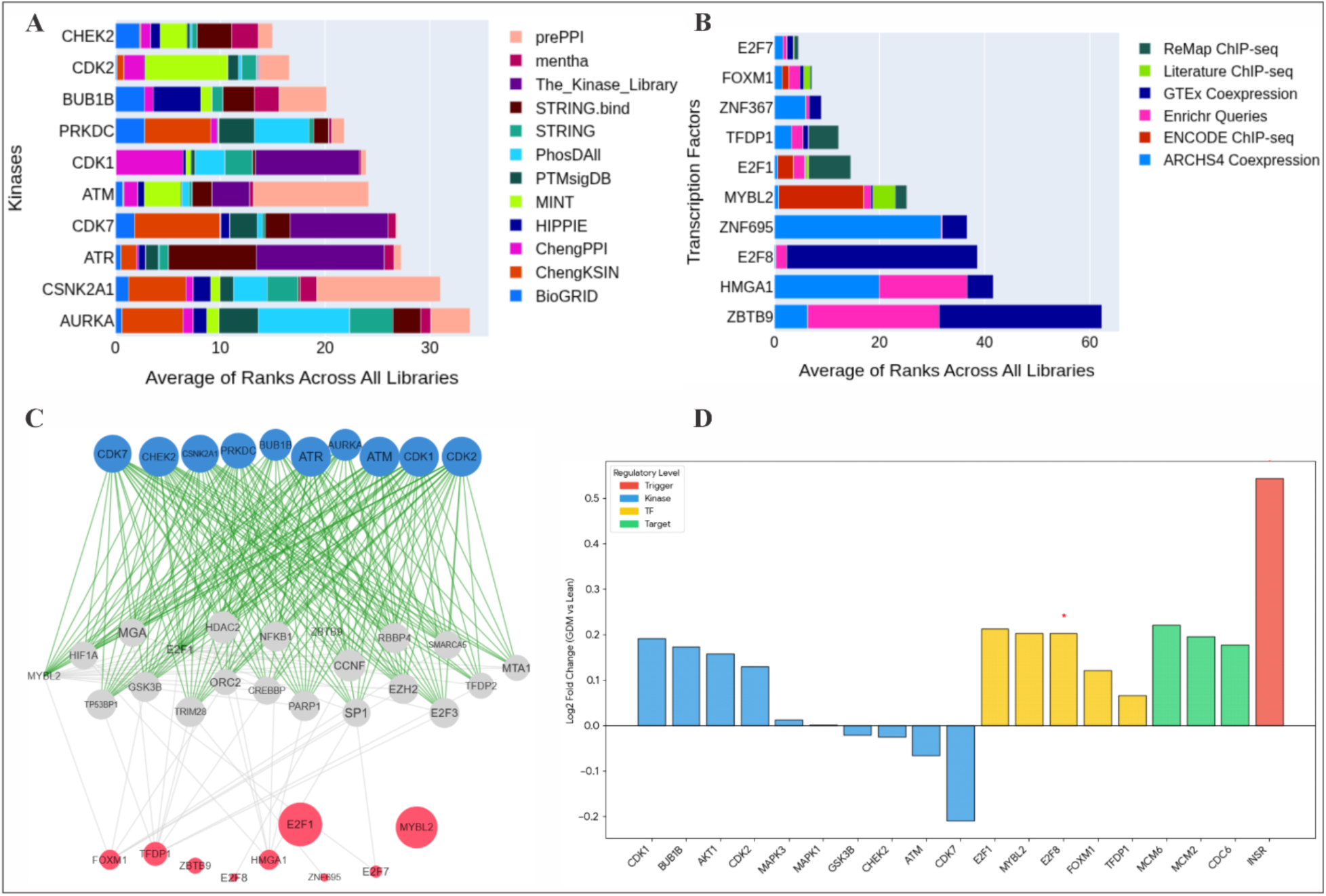
Integrated kinase-transcription factor regulatory architecture of female GDM. (A) Consensus Ranking of Upstream Kinases (KEA3). Top-ranked kinases inferred from the 16 female-specific leading-edge genes, with ranks averaged across experimental and predictive libraries (e.g., BioGRID, String). Primary signaling drivers include CHEK2, CDK2, and ATR, indicating a robust replication-stress response. (B) Consensus Ranking of Master Transcription Factors (ChEA3). Highest-ranked TFs coordinated by the upstream kinome, integrated across ChIP-seq (ReMap, ENCODE) and co-expression (ARCHS4) datasets. The dominance of E2F7/8 and FOXM1 corroborates motif-based findings of a sex-specific cell cycle re-entry axis. (C) Integrated X2K Regulatory Network. Signaling topology connecting the upstream kinome to the downstream transcriptome. Blue nodes (kinases, e.g., ATM/ATR) are linked via green edges (phosphorylation) to a central subnetwork of Red nodes (master TFs, e.g., E2F1, MYBL2). Grey/White nodes represent intermediate protein-protein interactors. This architecture mechanistically links GDM-induced metabolic stress to the proliferative transcriptomic signature unique to the female cohort. (D) Molecular Chain of Command. A cross-layered expression profile comparing the fold change of the metabolic trigger (*INSR*), upstream kinases (*CDK/BUB1B*), master transcription factors (*E2F/MYBL2*), and downstream target genes (*MCM/CDC*). Asterisks indicate significant differential expression in the female GDM vs. Lean comparison. The robust upregulation of downstream targets despite stable TF mRNA levels suggests a model of post-translational E2F1 activation in the female maternal-fetal interface.

**Fig 8:**
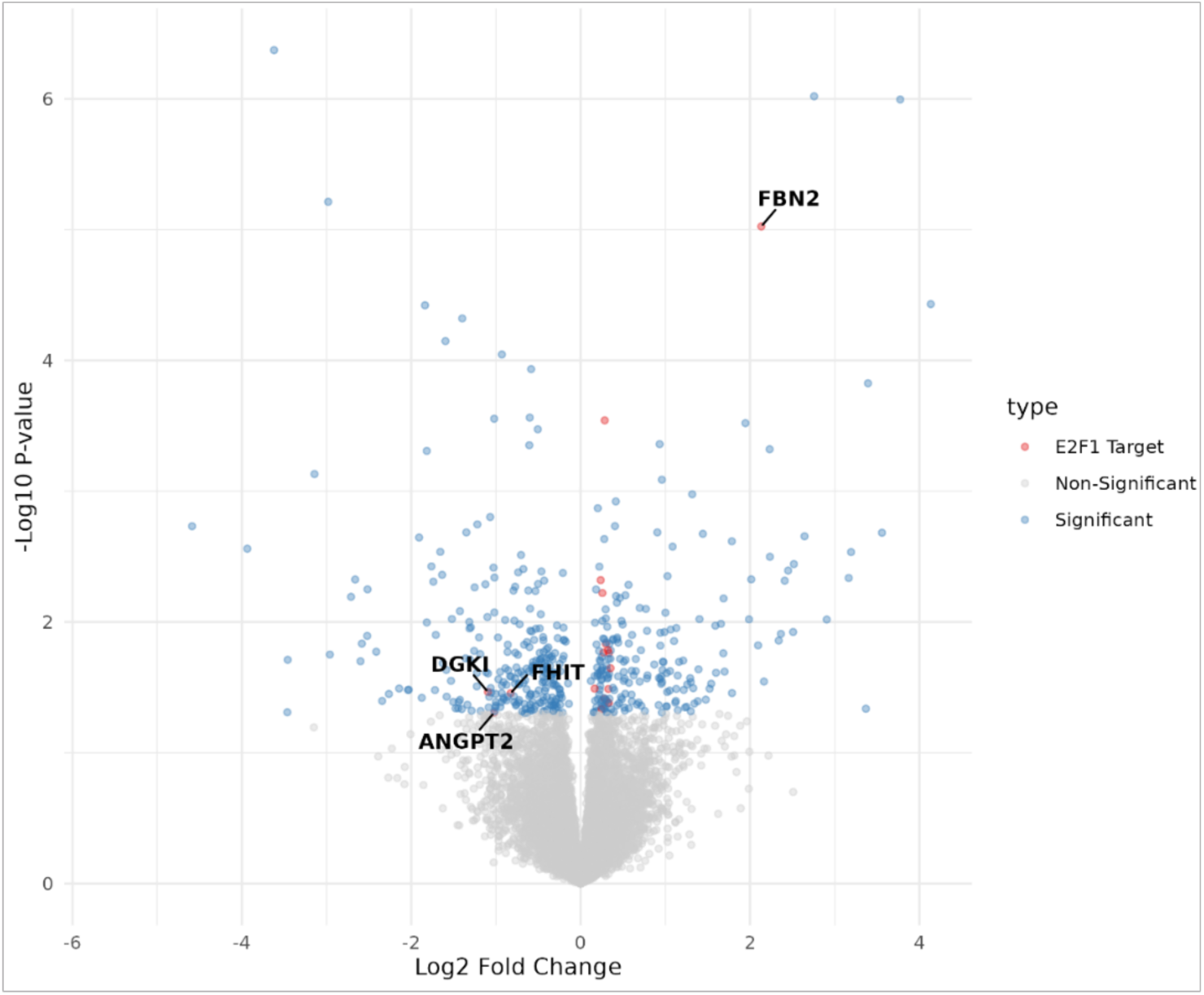
Global transcriptomic impact of the E2F1 Regulon. Volcano plot illustrating the differential gene expression landscape of the female GDM placenta compared to lean controls. Each point represents a single gene, plotted by its log_2_ fold change (x-axis) and statistical significance (−log_10_ p-value, y-axis). Blue nodes denote statistically significant differentially expressed genes (DEGs), while grey nodes represent non-significant transcripts. Red highlights identify high-confidence targets of the master regulator E2F1, as validated by the CollecTRI gene regulatory network. The clustering of E2F1 targets (e.g., FBN2, ANGPT2, FHIT, and DGKI) primarily within the significant quadrants—particularly among the upregulated features—confirms the functional dominance of the E2F1-driven proliferative program in the female GDM response.

### 3.7 Functional inference of transcription factor activity

To validate the upstream regulatory architecture and distinguish between transcript abundance and functional potency, we performed transcription factor (TF) activity inference using the *decoupleR* package. Activity scores were calculated via the Unweighted Least Squares (ULM) algorithm using the CollecTRI gene regulatory network. This approach identifies the functional status of a TF based on the coordinated expression of its downstream targets. The female GDM regulatory landscape was dominated by the robust functional activation of the *E2F* family, led by *E2F1* (*z* = 7.96, *p* =1.77×10^−15^), *E2F4* (*z* = 7.20), and *E2F3* (*z* = 5.98) (Table 2). This high positive *z*-score confirms that the *E2F* family is the primary functional engine driving the G1/S transition in the female placenta, despite the stable mRNA levels of the TFs themselves observed in the previous analysis. This proliferative drive was further supported by the significant activation of *MYC* (*z* = 5.63), a master regulator of metabolic reprogramming. Conversely, the analysis identified a systemic functional suppression of transcription factors essential for innate immunity and vascular maturation. The most significantly repressed regulator was *IRF1* (z = −5.67, p = 1.42×10^−8^), a key mediator of Type-I interferon signaling, followed by the NF-κꞵ subunit *REL* (z = −4.46) and *STAT1* (z = −4.21). Collectively, these activity scores demonstrate a functional polarization in the female GDM phenotype: a gain of mitotic function paired with a loss of immunovascular homeostatic control.

**Table 2:** Top 10 Inferred transcription factor activities in female GDM.

| Rank | TF | Activity score (z) | p-value | Activity status | Primary biological function |
| --- | --- | --- | --- | --- | --- |
| 1 | *E2F1* | 7.96 | $1.77e^{-15}$ | ↑ | DNA replication & S-phase entry |
| 2 | *E2F4* | 7.20 | $6.40e^{-13}$ | ↑ | G1/S transition & cell cycle control |
| 3 | *E2F3* | 5.98 | $2.28e^{-9}$ | ↑ | Regulation of cell proliferation |
| 4 | *E2F2* | 5.70 | $1.23e^{-8}$ | ↑ | Orchestration of cell cycle progression |
| 5 | *IRF1* | -5.67 | $1.42e^{-8}$ | ↓ | Innate immune response & IFN signaling |
| 6 | *MYC* | 5.63 | $1.82e^{-8}$ | ↑ | Metabolic reprogramming & biosynthesis |
| 7 | *FOXF1* | -5.20 | $1.97e^{-7}$ | ↓ | Angiogenesis & vascular development |
| 8 | *HIVEP2* | -4.81 | $1.97e^{-7}$ | ↓ | Inflammatory homeostasis |
| 9 | *REL* | -4.46 | $8.42e^{-6}$ | ↓ | NF-κB mediated immune survival |
| 10 | *STAT1* | -4.21 | $2.61e^{-5}$ | ↓ | Type-I/II interferon alarm response |

Note: ↑ mean up-regulated and ↓ downregulated

### 3.8 Regulatory landscape visualization

To visualize the regulatory reach of the identified master regulator, known *E2F1* targets were overlaid onto the global female GDM transcriptomic volcano plot (Figure 9). The distribution of these targets confirms that *E2F1* orchestrates a broad genomic reprogramming event rather than a unidirectional shift. While significant targets such as *FBN2* (*log_2_FC* > 2.0) were strongly upregulated, other key members of the regulon, including the angiogenic regulator *ANGPT2* and metabolic modulator *DGKI*, were significantly downregulated (*log_2_FC* < -1.0). This coordinated, multi-directional shift in validated targets—including the tumor suppressor *FHIT*—spatially demonstrates that the *E2F1* hub serves as a central architect of the female GDM response. The enrichment of these targets across both the activated and suppressed quadrants of the volcano plot provides evidence that *E2F1* activity is linked to the fundamental divergence of cellular resources, prioritizing a specific proliferative program while simultaneously modulating markers associated with vascular maintenance and metabolic signaling.

**Fig 9:**
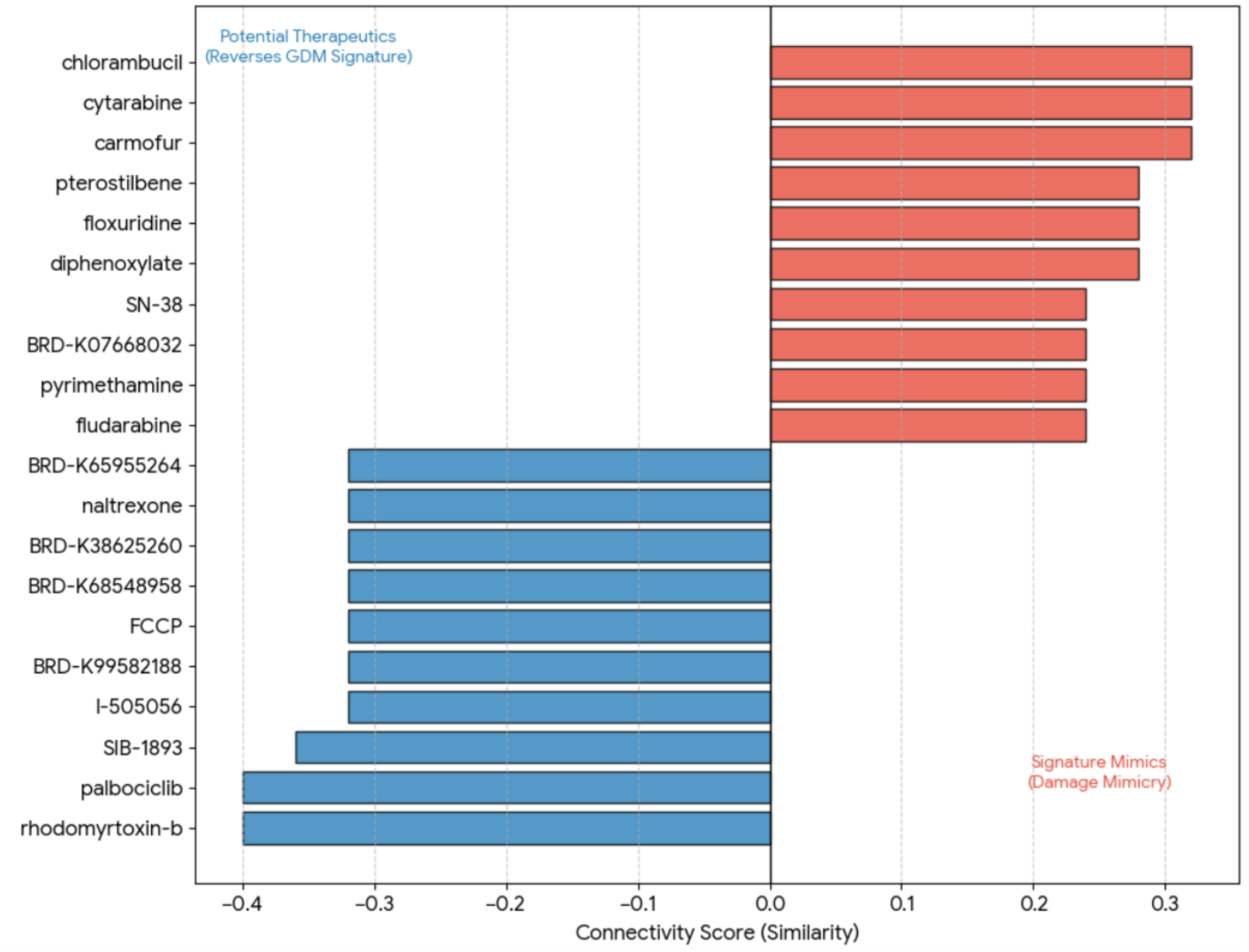
Connectivity mapping of GDM signature reversal and mimicry. A ranking of small molecules from the LINCS L1000 database based on their transcriptional similarity to the female GDM profile. Blue bars represent potential therapeutics exhibiting Signature Reversal (negative connectivity), led by the CDK inhibitor Palbociclib. Red bars represent Signature Mimics, primarily consisting of DNA-damaging agents. This divergence illustrates the transition from metabolic trigger to cellular stress and identifies specific pharmacological targets for mitigating GDM-induced placental damage.

### 3.9 Therapeutic connectivity: signature reversal and mimicry analysis

To translate the female-specific *E2F1*-centric regulatory architecture into therapeutic avenues, connectivity mapping was performed using the L1000FWD platform against the LINCS L1000 library. This analysis identified small molecules capable of either reversing the GDM transcriptomic signature (reversal agents) or mimicking the pathological profile (mimetic agents) (Figure 10). Palbociclib, a selective CDK4/6 inhibitor, emerged as a top candidate for signature reversal (Connectivity Score = -0.40). The pharmacological ability of a CDK inhibitor to negatively correlate with the GDM profile provides independent validation that the female pathology is driven by a hyper-active *CDK-E2F1* axis. Additional reversal candidates included the metabolic and mitochondrial modulators Naltrexone and FCCP, suggesting that the GDM phenotype involves a secondary component of mitochondrial stress. Conversely, connectivity analysis revealed that the GDM transcriptomic signature demonstrated significant positive similarity to several DNA-damaging agents and antineoplastics, including Chlorambucil, Cytarabine, and Carmofur (Similarity > 0.30). This chemo-mimetic footprint indicates that the female maternal-fetal interface in GDM exhibits a transcriptional state analogous to that induced by cytotoxic agents. Collectively, these pharmacological findings corroborate our mechanistic model: the female GDM response is defined by a state of replication stress and forced cell-cycle progression, which can be computationally reversed through targeted cell-cycle inhibition.

**Fig 10:**
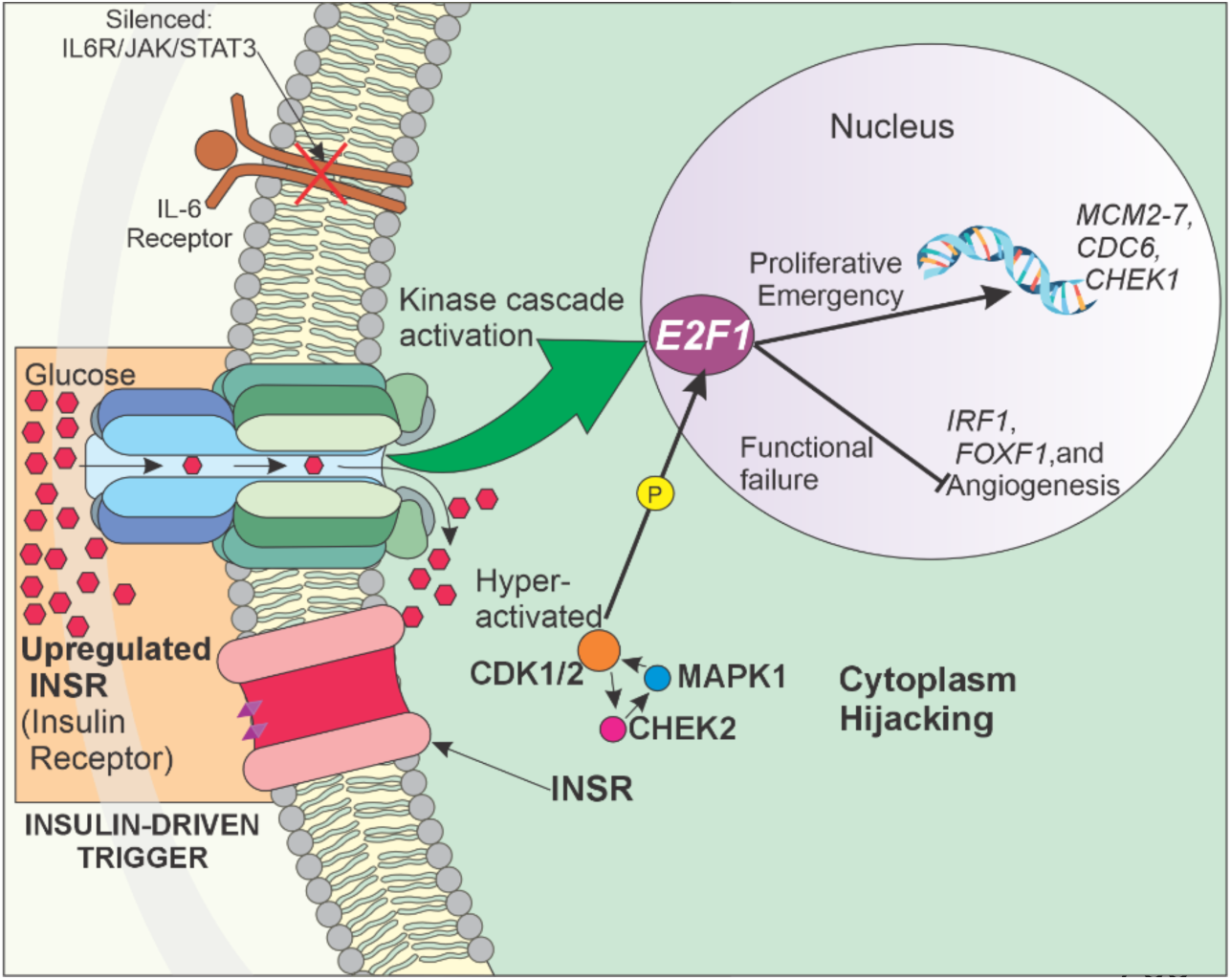
Proposed GDM-proliferative emergency: hijacking the cell cycle and silencing function

## Discussion

The findings of this study provide a high-resolution molecular map of how the intrauterine environment under GDM stress re-wires the fetal-placental endothelium. By shifting the unit of analysis to purified ECFCs, we have identified the specific transcriptional engines—the E2F1 and AR regulons—that drive sex-specific vascular programming. Our data suggests that GDM does not impose a monolithic metabolic burden; rather, it triggers two distinct biological trajectories. While the male placenta adopts a minimalist, AR-driven metabolic adjustment, the female placenta undergoes a massive, E2F1-driven proliferative emergency. This divergence provides a mechanistic basis for the fuel-mediated teratogenesis described by [2], suggesting that DOHaD-associated risks result from sex-specific failures in cellular homeostatic control.

Our analysis reveals that maternal obesity acts as a sexually dimorphic priming event that pre-configures the placental transcriptome. In males, this priming is predominantly metabolic, targeting insulin signaling pathways such as *AKT3*, aligning with evidence that male progenitors are significantly more sensitive to maternal metabolic shifts [6]. Conversely, the female placenta exhibits a unique structural and lipid-vascular priming. While we identified remodeling markers like *CORIN* and *FAP*, previous reports noted that maternal obesity is associated with higher cord blood triglycerides specifically in female fetuses [5]. This suggests that prior to a GDM diagnosis, the female interface is already managing a higher lipid load and a stiffened extracellular matrix (ECM) scaffold, creating a pre-stressed environment that sensitizes the placenta to subsequent GDM insults.

The RRHO analysis identifies a transcriptional collapse of the homeostatic niche unique to the female vasculature. While both sexes respond to GDM stress (indicated by the shared upregulated hotspot), the silencing of 3,004 genes in females—including core *TGF-ꞵ* and *JAK-STAT3* repair circuits—suggests a profound failure to maintain vascular identity. We propose that this shift creates a chemo-mimetic emergency where female cells abandon organized morphogenesis in favor of unscheduled mitogenesis. A central discovery of this study is the identification of E2F1 as the master regulator of this phenotype. While *E2F1* mRNA levels remained stable, its functional regulon—including *MCM2-7* and *CDC6*—showed unprecedented regulatory convergence (NES = 16.86). We validated this using transcription factor activity inference, confirming the female landscape is dominated by the robust functional activation of the E2F family: E2F1 (*z* = 7.96, *p* =1.77×10^−15^), *E2F4* (*z* = 7.20), and *E2F3* (*z* = 5.98) (Table 2). This provides high-confidence evidence that E2F1 is the primary functional engine driving the G1/S transition despite transcriptional stability.

We propose that this unshackling of E2F1 represents a failure of inhibition rather than a surge in synthesis (Figure 10). GDM-induced stress likely triggers a signaling relay—potentially involving the methylation of CDK inhibitor promoters such as *p21*(1) or a kinase cascade (CDK1/2, ATR) —that post-translationally activates E2F1. This drive is amplified by the activation of *MYC* (z = 5.63), a master regulator of metabolic reprogramming. Importantly, our results reveal a profound functional polarization: a gain of mitotic function paired with a systemic suppression of transcription factors essential for innate immunity and vascular maturation. The significant repression of *IRF1* (*z* = -5.67, p = 1.42 10^−8^), a key mediator of Type-I interferon signaling, followed by the NF-κꞵ subunit *REL* (*z* = -4.46) and *STAT1* (*z* = -4.21), demonstrates a complete loss of immunovascular homeostatic control.

This sex-specific divergence is corroborated by findings in fpECs where Hippo signaling was identified as a female-specific hit [4]. This provides a direct bridge to our E2F1 results; a Hippo-off state acts as a master trigger for the uncontrolled expansion of tissue-specific stem cells (30,31) The simultaneous activation of E2F1 and the suppression of *TGF-ꞵ* maintenance circuits suggest hierarchical reprogramming. Consistent with established crosstalk, the Hippo-YAP axis can antagonize Smad-mediated signaling and modulate *STAT3* responses. In the female GDM environment, the hijacked Hippo-E2F1 axis functionally overwrites morphogenic instructions, effectively silencing the maintenance crew to prioritize immediate, albeit dysfunctional, cellular expansion.

Ultimately, this hierarchical collapse suggests a fundamental difference in biological survival logic between the sexes. While males adopt a minimalist metabolic adjustment, the female vasculature engages in a systemic transcriptional retreat. By leveraging the 14q32 miRNA cluster [5]—a master regulator of neovascularization (4)—the female progenitor niche prioritizes immediate survival through rapid mitotic expansion over long-term structural integrity. This retreat has immediate functional consequences: the reduction of key morphogenic regulators like *lncRNA KLRK1-AS1*, which significantly delays wound healing in neonatal endothelial cells (7) directly connects our molecular observations to the physical reality of delayed villous maturation seen clinically.

However, this strategy is unsustainable. As defined by the hallmarks of cellular aging [32] forced, unscheduled proliferation leads to regenerative exhaustion. This bioinformatic model explains the physical hyper-proliferation observed by [33] and the structural failures reported by [34]. True ECFCs must possess robust clonal proliferative potential;(35) however, the E2F1-driven surge forces these progenitors into an emergency state that prematurely exhausts this potential. This exhaustion likely transitions the cells into immunomodulatory endothelial cells (IMECs) [36] where they abandon their role as vascular builders to become immune sentinels—marked by the upregulation of HLA-class markers and immune-trafficking genes.

Our findings provide a mechanistic basis for the tissue-level dimorphism recently reported by [5]. While they observed female-specific enrichment in immunoregulatory pathways and altered *IGFBP1* expression in bulk placental tissue, our model demonstrates that these are likely the downstream consequences of the progenitor-level proliferative emergency identified here. Despite the mechanistic clarity provided by our sex-stratified approach, we acknowledge that the feto-placental response may be further modulated by maternal ancestry and racial background. Population-level data indicate that GDM prevalence and its associated cardiovascular sequelae disproportionately affect specific ethnic cohorts [37,38]. However, our current analysis was limited by the demographic distribution of the primary dataset. Future studies utilizing larger, multi-ethnic cohorts are required to determine if the E2F1-driven proliferative emergency is intensified by specific racial epigenetic predispositions, which would allow for even more refined risk-stratification models.

The clinical significance of this model is punctuated by our small-molecule reversal analysis. The discovery that the female GDM signature mimics the effects of genotoxic agents suggest the placenta perceives high glucose as a DNA-damaging insult. The emergence of Palbociclib (a CDK4/6 inhibitor) as a top reversal candidate confirms that targeted cell-cycle arrest can restore homeostatic balance. As NIPT makes fetal sex determination routine, our findings provide a framework for sex-specific vasculo-protective interventions that move beyond monolithic glucose management.

## Conclusion

Our study identifies a fundamental functional asymmetry in how the fetal-placental vasculature responds to GDM, demonstrating that the molecular scars of intrauterine stress are not just different in degree between the sexes, but fundamentally different in kind. While the male interface exhibits metabolic resilience through minimalist adjustments to manage maternal glucose spikes, the female cohort undergoes a maladaptive, E2F1-driven proliferative emergency. By shifting the unit of analysis to the integrated Kinase-TF-Pathway axis, we have characterized this female-specific response as a double-insult trajectory culminating in replicative exhaustion. This process begins with an obesity-driven priming phase that imposes a female-specific lipid load and pre-configures the extracellular matrix scaffold. The subsequent insult of chronic GDM hyperglycemia subsequently uncouples mitogenesis from structural morphogenesis, post-translationally activating the E2F1 engine through the disruption of upstream kinase signaling and the suppression of cyclin-dependent kinase inhibitors.

Ultimately, this pathway activation forces a systemic transcriptional retreat. To prioritize immediate cellular survival, female progenitors compromise long-term structural and immunovascular integrity—marked by the loss of critical morphogenic regulators like *lncRNA KLRK1-AS1*, the silencing of core *TGF-ꞵ* and *JAK-STAT3* repair circuits, and the suppression of innate immune networks (*IRF1*, *STAT1*). This hierarchical collapse drives a profound phenotypic transformation, forcing true endothelial colony-forming cells to abandon their roles as vascular builders and adopt an exhausted, immune-transformed sentinel state. Clinically, this bioinformatic model provides a precise molecular explanation for the physical hyper-proliferation and delayed villous maturation observed in GDM pregnancies.

In an era of accelerating precision medicine and routine non-invasive prenatal testing (NIPT), these findings provide a solid mechanistic framework for identifying the cell cycle as a primary, sex-specific therapeutic target. Our results advocate for a vital paradigm shift toward sex-stratified obstetric care and targeted vasculo-protective interventions, disrupting the developmental programming of adult cardiovascular disease before birth to protect the health of the next generation.

## Ethics approval and consent to participate

Not applicable

## Consent for publication

Not applicable

## Availability of data and materials

The raw transcriptomic data analyzed in this study are available in the National Center for Biotechnology Information (NCBI) Sequence Read Archive (SRA) database under BioProject accession number PRJNA885556. These data were originally generated and described by (Weiss et al., 2023) in their investigation of gestational weight gain and fetal endothelial function. The processed data and custom bioinformatic scripts used in this study are available from the corresponding author upon request.

## Competing interests

The authors declare there had no competing interests

## Funding

This research was supported by a USDA-NIFA research grant 2023-67016-39917 (BNT). The funders had no role in study design, data collection and analysis, decision to publish, or preparation of the manuscript.

## Authors’ contributions

MSA, OB and BNT conceived the idea; MSA and OB carried out the study; MSA and OB analyzed the data and wrote the initial draft; MSA, OB, OBM, OO and BNT contributed to the scientific content and final work. All authors approved the final work for publication.

## Acknowledgments

This work constitutes an ongoing collaboration between Systems Immunology and Computational Biology Research group, Rochester Institute of Technology and the Center for Emerging and Re-emerging Infectious Diseases, Ladoke Akintola University of Technology, Ogbomosho, Nigeria. Part of this work was presented at the American Association of Immunologists Annual Conference, Boston MA; April 15–19, 2026. We are grateful to the College of Health Sciences and Technology for ongoing support. The authors acknowledge the Rochester Institute of Technology Research Computing for providing the high-performance computing (HPC) resources used for the downloads of SRA data and transcriptomic analysis.

## Notes

### Competing Interest Statement

The authors have declared no competing interest.

